# Comparison of the structural organization of *reeler* hippocampal CA1 region with wild type CA1

**DOI:** 10.1101/508648

**Authors:** Malikmohamed Yousuf, Shanting Zhao, Michael Frotscher

## Abstract

The dendritic pattern defines the input capacity of a neuron. Existing methods such as Golgi impregnation or intracellular staining only label a small number of neurons. By using high-resolution imaging and 3D reconstruction of green fluorescent protein-expressing neurons, the present study provides an approach to investigate the anatomical organization of dendritic structures in defined brain regions. We characterized the structural organization of dendrites in the CA1 region of the mouse hippocampus by analyzing Sholl intersections, dendritic branches, branching and orientation angles of dendrites, and the different types of spine on the dendritic branches. Utilizing this quantitative imaging approach, we show that there are differences in the number of Sholl intersections and in the orientation of apical and basal dendrites of CA1 pyramidal neurons. Performing 3D reconstructions of the CA1 region of the *reeler* hippocampus, we show that neurons of this mutant display an arbitrary orientation of apical dendrites at angles ranging from -180 to +180 degrees in contrast to wild-type mice that show a preferred orientation angle. This methodology provides a way of analyzing network organization in wild type and mutant brains using quantitative imaging techniques. Here, we have provided evidence that in reeler a sparse, weakly connected network results from the altered lamination of CA1 pyramidal neurons and the variable orientation of their dendrites.

**Highlights:**

- High-resolution imaging and 3D reconstruction of CA1 pyramidal neurons
- Analysis of Sholl intersections
- Analysis of the orientation of apical and basal dendrites of CA1 pyramidal neurons
- Altered dendrite orientation angle of apical dendrites in *reeler* mutant mice
- More stubby spines compared to thin and mushroom spines in wild-type CA1

## 1. Introduction

Our knowledge about the structure, function and connectivity patterns of the hippocampus and its associated structures dates back to the classical Golgi studies by Golgi (1896), Ramón y Cajal (1952) and Lorente de Nó (1934). The integration of synaptic inputs by a neuron and consequent action potential generation depends on many factors. One of the key features controlling this process is the complexity of the dendrite branches and their length (Vetter et al., 2001). Dendrite branching pattern and the dendritic field are involved in the normal functioning of many physiological processes. The area covered by dendritic branches influences the extent of the inputs the neuron can receive and compute, while the complexity of dendritic branching determines its specialized task (Agmon-Snir et al., 1998; Borst et al., 2010; Euler et al., 2002; Single and Borst, 1998). The geometry of dendritic branches, the number of synaptic sites, the distribution of different types of spines on the dendritic branches, the distribution of different voltage-gated ion channels, and the history of previous synaptic activity are all factors that determine how the dendrites integrate the incoming information that the cell receives from multiple inputs spanning across the dendritic branches. Moreover, the computation of the proximal dendrites differs from that of the distal branches, and the summation of their response also varies (Losonczy and Magee, 2006; Polsky et al., 2004). For instance, the attenuation of EPSPs (Excitatory Post-Synaptic Potentials) is steeper in basal dendrites than in apical dendrites when they propagate to the soma (Inoue et al., 2001). Besides that, basal dendrites display significantly attenuated back-propagating action potentials (APs), which suggests that basal dendrites integrate information in a sub-threshold way. The mode of input summation in basal dendrites is location-dependent, in contrast to the spatial amplification of temporally clustered inputs processed by apical or distal dendrites (Nevian et al., 2007). To generate action potentials and to surmount the passive properties of dendrites, neurons use summation of synaptic potentials either spatially or temporally, or in a combination of both. When co-incident inputs are impinging at different locations within the branch or in the tree, the depolarization caused by the concomitant input will not dampen the driving force of other synaptic inputs. Consistently, co-incident inputs impinging spatially or temporally across the branches or in the tree (Markram et al., 1997; Williams and Stuart, 2003) create different summations, which may be sublinear, linear or supralinear. The morphology of the dendritic tree plays a pivotal role as the increase in the number of branch points (Vetter et al., 2001) attenuates the EPSPs that reach the soma, which dampens the generation of an AP. It is important to consider that synaptic events are changes in membrane conductance, and the shapes of the responses are influenced by the geometry or the morphology of the dendritic branching architecture. A study combining experimental data and computational simulations (Vetter et al., 2001) showed how the dendritic morphology and geometry play a pivotal role in governing the forward and backward propagation of APs. The authors demonstrated that the number of dendritic branching points determine the efficacy of propagation. The more dendritic branches, the greater the attenuation of charge. Hence, the propagation of an AP is dampened both forward to the soma and backward to the distal dendrites. This partially explains why Purkinje neurons with a large dendritic geometry are incapable of propagating the synaptic potentials to the distal regions. In summary, the morphological and the geometrical parameters are one of the key features that determine the efficacy of propagation to dendrites. Unless there is a compensated increase in Na^+^ channel density, the increase in the number of dendritic branches will attenuate the back propagation of APs into the dendrites. An increase in complexity should also be compensated by an increase in the number of synaptic inputs so that there is enough depolarization to overcome the active and passive properties of the dendrites to reach the distal portions so that they can act as a coincidence detector for strengthening those synapses that are activated. In addition, the branching pattern and orientation of dendrites are important since the orientation of dendrites determines how they become a distributed circuit (Jia et al., 2010; Varga et al., 2011). The spines present on the dendrites enable an increase in connectivity with the bypassing axons allowing them to maximize their input.

In the present study, we aim to unravel the structural organization of the dendrites in the CA1 region of the hippocampus. We reconstructed the dendrites emanating from the somata of individual pyramidal neurons, which allowed us to characterize the number of dendritic intersections and dendritic branches, which directly describes the dendritic complexity, and indirectly, the capabilities of the network which the characterized neurons are part of. To highlight the usefulness of this approach, we decided to analyze the structural organization of apical dendrites in the CA1 region of *reeler* mice. Reelin is a heavily glycosylated extracellular matrix protein with a molecular weight of 400 kDa (D’Arcangelo et al., 1999; D’Arcangelo et al., 1995). In humans, abnormal migration of neurons leads to lissencephaly (Walsh, 1999), and one type of lissencephaly is caused by a defective *reelin* gene (Hong et al., 2000). Reelin is expressed by GABAergic interneurons in the stratum radiatum and stratum oriens of the CA1 region in the adult hippocampus (Campo et al., 2009). Histological work by Stanfield and Cowan (1979a; 1979b) has shown that the apical dendrites in the deep pyramidal layer of the CA1 region of the *reeler* hippocampus are inverted and mis-oriented. However, high-resolution studies aimed at understanding the structural and the network organization of the CA1 region of the hippocampus in *reeler* mice are still lacking. By analyzing the structural organization of apical dendrites in *reeler*, we were able to determine structural changes in comparison to wild-type neurons, thus providing insights into the organization of the neuronal network and the connectivity matrix in the reeler mutant. Hypotheses put forward in light of the present results warrant further electrophysiological experiments to corroborate our morphological findings. This would eventually help to dissect the computational capability of the network each individual neuron is part of in the *reeler* mutant hippocampus.

## 2. Materials and Methods

### 2.1. Experimental animals

Three adult wild-type mice expressing enhanced green fluorescent protein (eGFP) under the Thy1 promoter (C57BL/6J, Thy1-eGFP-M, Jackson Laboratory; Feng et al., 2000) and 3 adult *reeler* mutants (C57BL/6J, Jackson Laboratory; Falconer, 1951) also expressing eGFP under the Thy1 promoter were used for the present study. Animals were reared in the animal facility under normal conditions (22 – 24° C and 55% humidity). Animal care and handling of the animals before and during the experiments was in accordance with the European Union regulations.

### 2.2. Tissue preparation for light microscopy studies

Animals were anesthetized deeply by Narcodorm-n (pentobarbital, 180mg/kg, i.p., Alvetra, Germany). They were then perfused transcardially with 0.9% saline (NaCl) for 1 minute followed by 4% paraformaldehyde (PB, Merck, Germany) in 0.1 M phosphate buffer (PFA, Merck, Germany), for 15 minutes. Brains were removed and post-fixed overnight. They were cut horizontally or coronally on a vibratome (VT1000S, Leica, Germany) at a thickness of 50 μm. Sections were embedded with DAKO mounting medium (DAKO).

### 2.3. Imaging the hippocampal slices with a custom made macro plug-in

The fixed hippocampal slices were imaged using a custom based macro plugin (Dr.Jin, Life Imaging Center, Zentrum for Biosystem analysis, personal communication). To image the whole CA1 of the hippocampus of the wild type and *reeler*, the plugin was used, enabling the user to image the whole structure in to an array of tiles with an overlap of 10%. This allowed the user to stitch the tiles using commercially available programs such as Xuv stitch tools (http://www.xuvtools.org/), rendering the reconstruction in 3 dimensions. The macro was designed in a way that it can scan long duration experiments covering different scan areas. The hippocampus is broadly distributed into three different scan areas: the dentate gyrus (including the granule cell layer and the outer molecular layer), the CA3 and the CA1. Each of the areas has been scanned and imaged as an array of tiles with different scan settings such as pinhole size, bit depth, scan speed etc. Once the scan settings were assigned for the different scan areas, the designations are uploaded and are fully controlled by the macro. Broadly, the setup which enabled the macro to automatically image the whole structure in 3D (for instance CA1 of hippocampus) is divided in to 4 steps. First, the stage tilt was adjusted which compensated for the changes in the focus, when the stage was moved across different tiles during imaging. This step was pivotal as it enabled the user to image the whole structure without losing focus and it corrects any drifts in the movement of the stage. This was performed by measuring the positions of x, y co-ordinates of the four ends of the cover slip in relation to the position in the center using an autofocus mode. Once the stage tilt was adjusted, the focus of different tiles in the array was corrected to the x, y co-ordinates by the stage tilt test, while the stage was moved across individual tiles or across the array of tiles. It has to be taken into consideration that the sample should not be moved from its relative position once the stage tilt was measured and corrected.

The next key step was to create the overview of the structure of interest to be imaged. Once the overview was created, it was better to synchronize the image with a reference so that one could re-do or re-load the same image region for imaging the same structure in the future. The overview could be imaged in a single channel or in a multiple channel image plane of the specimen. Additionally, the overview enabled a microscope independent co-ordinate system. By synchronizing the overview, the hardware automatically synchronized with the image co-ordinates. For instance, in this case, we imaged the overview of the hippocampus comprising of the major three regions described above with a low resolution lens (5 – 10X) to define the structure. This overview structure was synchronized so that the x, y co-ordinates that relate to the center and the borders are marked rendering them for future uses. For further verification, one could check the co-ordinates of the overview structure by clicking the mouse on the overview and displaying the current co-ordinates with respect to the origin. Once the overview of the hippocampus was created, the scan settings were configured to scan-in the overview. Thy1::GFP hippocampal slices were excited with a laser having excitation wavelength of 488nm and the emitted light was collected at the emission wavelength of 513nm using the broad pass filter of 505 – 530. The only channel used was the GFP emission channel for collecting the emission light. Once the light path, laser, and channels were set, the whole slice was imaged in a low resolution with a fast scan speed so that the whole hippocampus as well as specific regions could be visualized very clearly to set up arrays of tiles in the region of interest for high resolution imaging of the structures in 3D.

The experimental plan for the region of interest was assigned in the plug-in; in this case, the CA1 of the hippocampus. The tile scan for a single marked tile in CA1 was assigned on the overview and the scan settings were configured and stored. The neurons in the slices were visualized and imaged using a 40x oil immersion lens (NA 1.3) on an LSM 510(Zeiss) inverted microscope. The mercury HBO lamp with a GFP filter was used for the primary visualization of neurons in the slices. *Thy1*::GFP hippocampal slices were excited by excitation wavelength of 488nm and the emitted light was collected at the emission wavelength of 513nm. The cover slip position was considered 0.00 micron and the imaging direction was upward. The lens immersion refractive index and the medium refractive index were matched (1.515). The dimensions of individual tiles (X, Y and Z) were of 223.89μm, 223.89μm and Z varied from 40 - 50μm. The voxel size of an individual tile (X, Y and Z) was 0.109μm, 0.109μm and the Z axis ranged from 0.4 – 0.6 μm. The power of the laser (488nm) used for excitation was always kept constant at 4%. Each line scanned across the whole image was averaged twice for all the imaged slices and the pixel dwelling was 1.60μs. The master (voltage) gain was varied according to the experimental set up and the digital gain was always kept constant at 1. The pinhole size was normally kept as a default value of 1 airy unit. The numbers of tiles encompassing the whole region or the region of interest in the CA1 were configured with the same settings were marked and the configuration was stored in the settings. Thus the scan tile group was defined to image multiple tiles. For 3D reconstruction, all the neighboring tiles encompassing the CA1 region of hippocampus had an overlap of 5 -10% and the zoom factor was kept always at 1 to eschew border artifacts that would render the stitching difficult. The experimental settings for the individual scan tile group were re-checked and the experiment was started. The whole experimental scan area was controlled by the assigned function of the macro. Once the experiment was finished, the tiles span the region of interest, in this case the CA1 region of the hippocampus of wild type and *reeler*. This would be subsequently processed by the data processing plug-in of the macro to have a position of the x,y, z co-ordinates of the tiles so that one could use the profile to stitch the tiles in 3D using a software, Xuv stitch tools (Emmenlauer et al., 2009) for the 3D re-construction.

### 2.4. Stitching the dataset (Tiles) of the CA1 region of the hippocampus for 3D reconstructions

Stitching is a form of image registration, which is a process of intra – subject rigid registration. The images (tiles) were acquired in a regular grid. The CA1 region of the hippocampus was stitched using Xuv Stitch tools (Emmenlauer et al., 2009). The profiles of the co-ordinates of the x, y axis of each overlapping tile were made by the plug-in of the macro (Dr.Jin, Zentrum fur Biosystem Analyze, Freiburg, Germany, personal communication). This profile enabled the stitcher tool to use the original co-ordinates generated by the macro rather than performing an exhaustive search of the overlapping tiles, which would render inconclusive representations of the final 3D image for reconstruction. Once the tiles representing the whole CA1 of the hippocampus were stitched, they were further processed for 3D reconstruction in Imaris (http://www.bitplane.com/go/products/imaris).

The filament tracer obtained by Imaris provided a unique opportunity to reconstruct the dendrites emanating from the soma of the pyramidal neurons. The somas of the neurons are segmented by means of intensity thresholding as the cell bodies show more contrast than their processes. To minimize tracing errors, we always used median filtering to reduce the noise in the images after stitching all the images prior to reconstructions. The soma of all the neurons in the CA1 region is marked manually and by doing this, we made sure that there was no error in detecting the soma of the neurons. Before assigning the diameter of the cell body and the dendritic branches, a random set of neurons were taken and the diameters were measured manually to make sure the threshold values for segmenting the soma and the dendritic branches were in the correct range. The seed point (the path loop) to detect the dendrites is used as a path point and Imaris uses the next path to the next seed point along the brightest route (highest voxel intensities). By this, dark areas will be avoided and cross connections to other cells are usually not made. The seed point itself is detected with a blob detector. A fast matching algorithm was used to connect the seed points, starting at the end of the dendrites working the way back to the starting point along the highest intensities. The algorithms used in filament tracing to detect the dendrites can be found on the resource page of bitplane website, http://www.bitplane.com/go/support/resources under the filament tracer label. To validate the approach, E-GFP microinjected single pyramidal neurons were reconstructed and the statistics of Sholl intersections, dendritic branches, dendritic branching angle, and orientation angles were extracted from individual neurons and they were cross compared to the data extracted from the large pool of reconstructed pyramidal neurons using filament tracer of Imaris.

Dendritic branching angle was defined as the angle between the extending lines from the branch point and its peripheral neighbor. Angles were marked with dashed blue arcs, α. (Picture adapted from Imaris reference manual).

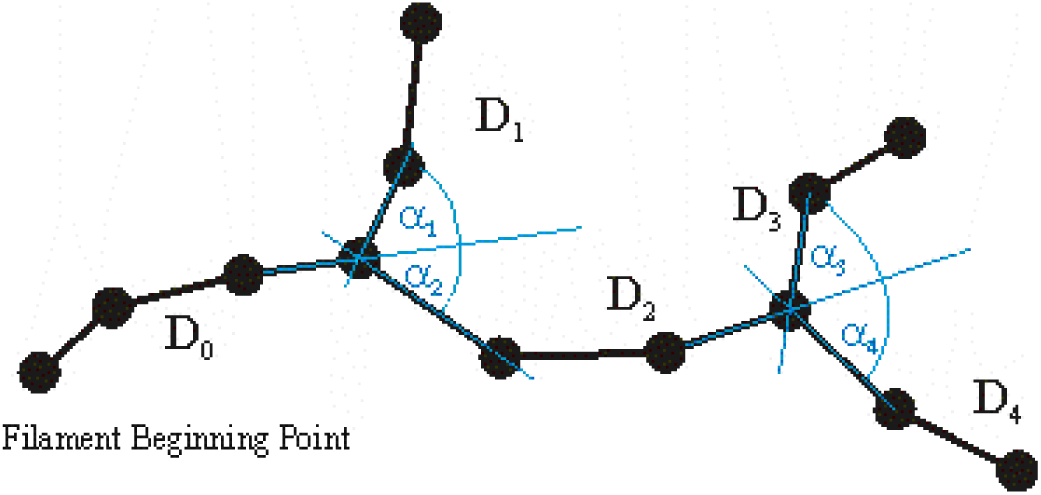

Dendrite orientation angle is defined as the angle formed between extending a line connecting distal vertices of the dendrite segment and the X-axis of the image within the XY plane. Angles were marked with dashed orange arcs, γ (Figure adapted from Imaris reference manual).

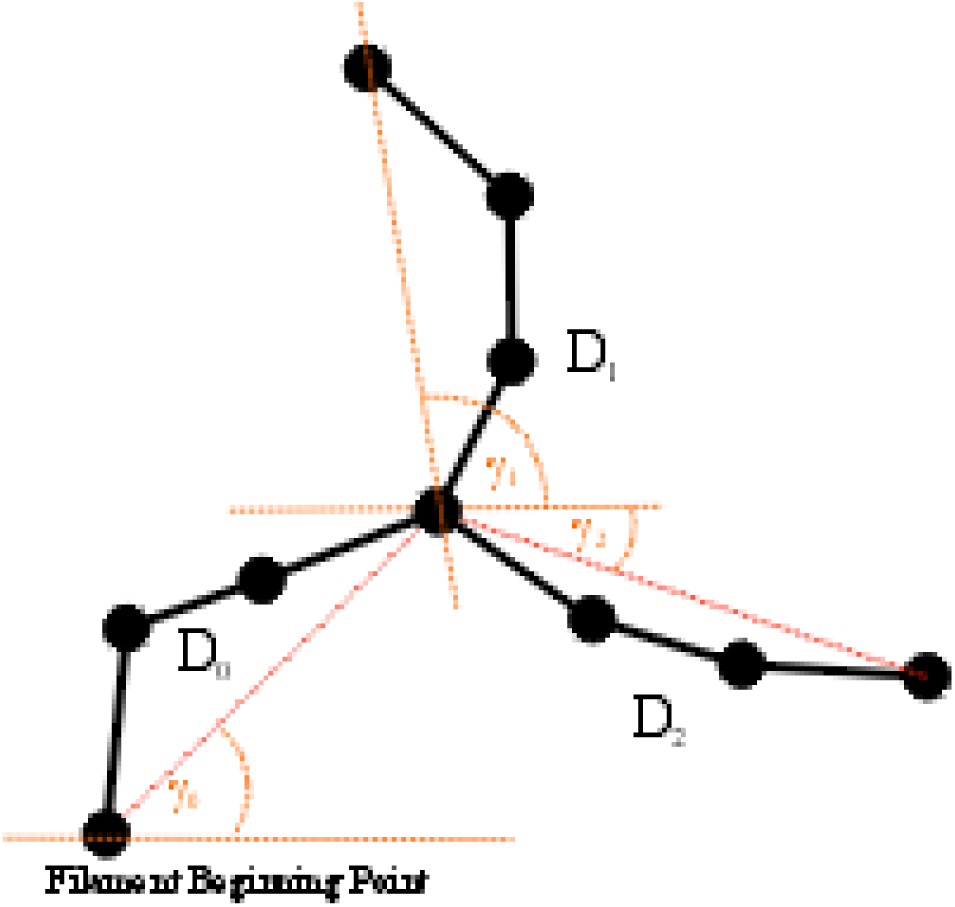

Sholl intersections were defined as the number of dendrite intersections (branches) on concentric spheres, describing dendrite spatial distribution as a function of distance from the beginning point (Sholl, 1953) The number of dendritic branches is calculated as the number of parallel-outstretched dendrite branches counted from the beginning point toward the terminal points at constant length increments.

### 2.5. Morphometric Analysis and Classification of Spines

The Morphometric analysis and classification of spines into different groups (Thin, Stubby, and Mushroom) was done in Neuron Studio (http://research.mssm.edu/cnic/tools-ns.html). The spine classification in the CA1 region of the hippocampus of wild type and *reeler* was done according to the instruction given by the program “Neuron Studio” with reference to the published work (Dumitriu et al., 2011). The deconvolution of all the images for classifying spines was done in Huygens’s Essential by Scientific Volume Imaging (http://www.svi.nl/HuygensDeconvolution). All the stacks of CA1 region in wild type and *reeler* were deconvolved with the same settings before they were processed for further analysis. The Z-smear in the image was corrected.

### 2.6. Statistical analysis

One-way ANOVA was used to test statistical differences of means between independent groups within the same experiment.

## 3. Results

### 3.1. 3D reconstruction of the pyramidal neurons in the whole CA1 of hippocampus provided a unique opportunity to dissect their structural properties of the apical and basal dendrites

In order to delineate the number of dendritic branches, branching pattern, orientation of dendritic angles and the Sholl intersections of the pyramidal neurons in the CA1 region, the whole CA1 of wild type mice was imaged and reconstructed in 3D. The array of tiles was stitched for the 3D reconstruction to delineate the structural changes in the dendritic tree as well as the spine morphology in the CA1 region of the hippocampus. In wild-type mice, pyramidal neurons are organized as a compact layer and their apical and distal dendrites are located in the stratum radiatum and lacunosum-moleculare, respectively, while the basal dendrites occupy the stratum oriens of CA1 (Fig. 1). Single pyramidal neurons were reconstructed before we went on to reconstruct the whole CA1 of hippocampus region. The 3D reconstruction now enables us to compare structural changes in wild type to that of reeler mice. For this endeavor, we reconstructed the whole CA1 region of reeler mice. Contrary to wild type CA1 (Fig.1), *reeler* mice showed no characteristic laminar structure and their pyramidal layer was split into two layers, a deep and superficial pyramidal layer in the CA1 region (Stanfield and Cowan, 1979a, 1979b) as confirmed by our imaging experiments (Supplementary Fig. 1). Of note, *Thy1*-GFP expression was not observed in interneurons (Figure. 1). Therefore, the present approach of using GFP-stained cells for analyzing projection neurons was advantageous compared to Golgi impregnation since some of the misoriented, multipolar CA1 pyramidal cells in the *reeler* mutant were not easily differentiated from interneurons (Stanfield and Cowan, 1979a., b;). The dendrite orientation of the apical dendrites in the deep pyramidal layer was inverted and did not form a compact layer (Stanfield and Cowan, 1979a; 1979b). The superficial layer is considered to be more normal than the deep one. We could confirm these findings by high resolution imaging of the whole structure of CA1 both in wild-type and *reeler* mice (Fig. 1 and Supplementary Fig 1). We measured by Sholl analysis the extent of dendritic complexity and dendritic arborization. In Sholl analysis, a series of concentric spheres are drawn, centered at the target neuron’s soma. The number of dendrite intersections from the target neuron (branches) with the spheres is then counted, allowing the experimenter to measure the spatial distribution of dendrites as a function of distance from the soma. The changes in the dendritic arborization and structure determine the network activity ((Segev and London, 2000). The average number of intersections of apical dendrites at a distance of 50 μm from the soma in wild-type mice was 5.79 ± 0.19 (mean ± SEM), Fig.2A, while the average number of apical dendritic branches as a function of distance (50 μm) from the soma was 8.41 ± 0.35 (mean ± SEM), Fig.2B. The number of dendritic branches and intersections, dendritic geometry plays a pivotal role in input integration. To characterize this, we examined the distribution pattern of the branching angle of dendrites of pyramidal neurons (Fig. 2C). Dendritic branching angle was defined as the angle between the extending lines from the branch point and its peripheral neighbor for each individual pyramidal neuron. To understand the orientation of dendrites emanating from the soma of pyramidal neurons in CA1, we analyzed their orientation angle. Dendritic orientation was defined as the angle formed by extending a line connecting distal vertices of the dendrite segment and the x-axis of image within the x-y plane. Pyramidal neurons showed a preferential orientation of their apical dendrites on a range of certain degrees with respect to the X-axis of the image. In wild-type mice 53.95% of apical dendrites exhibited an orientation preference for the angles ranging from -40 to +40 degrees, with the other apical dendrites distributed at the remaining angles. To measure the probability distribution of the orientation of apical dendrites in detail, a heat map with a bin size of 2.8 degrees was generated (Fig. 2D).

**Figure 1.**
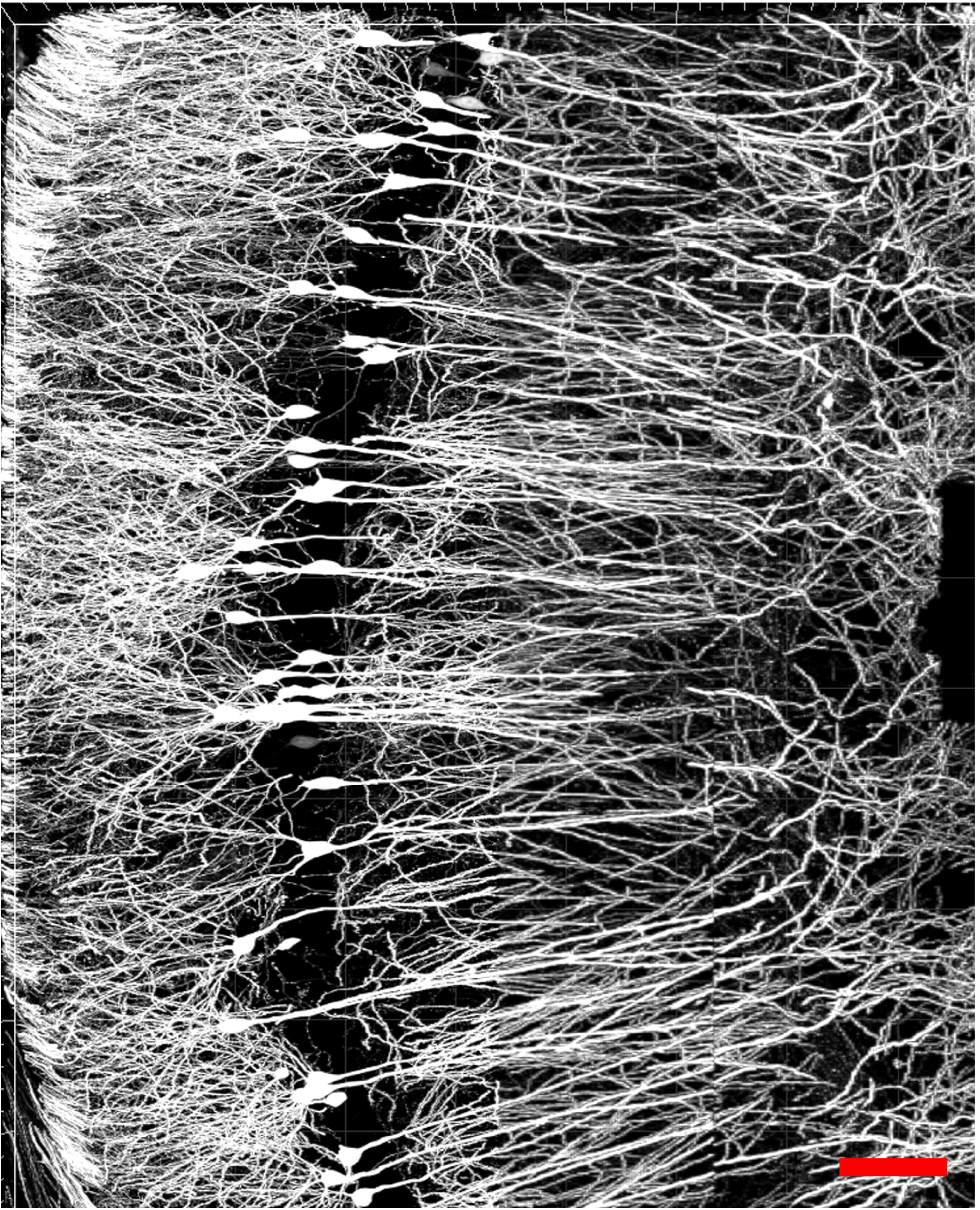
Reconstruction of the CA1 region of wild-type hippocampus. The CA1 region of wild-type hippocampus was documented by stitching 25 tiles with Xuv stitch tools and reconstructed using Imaris. Scale bar 100 μm.

**Figure 2.**
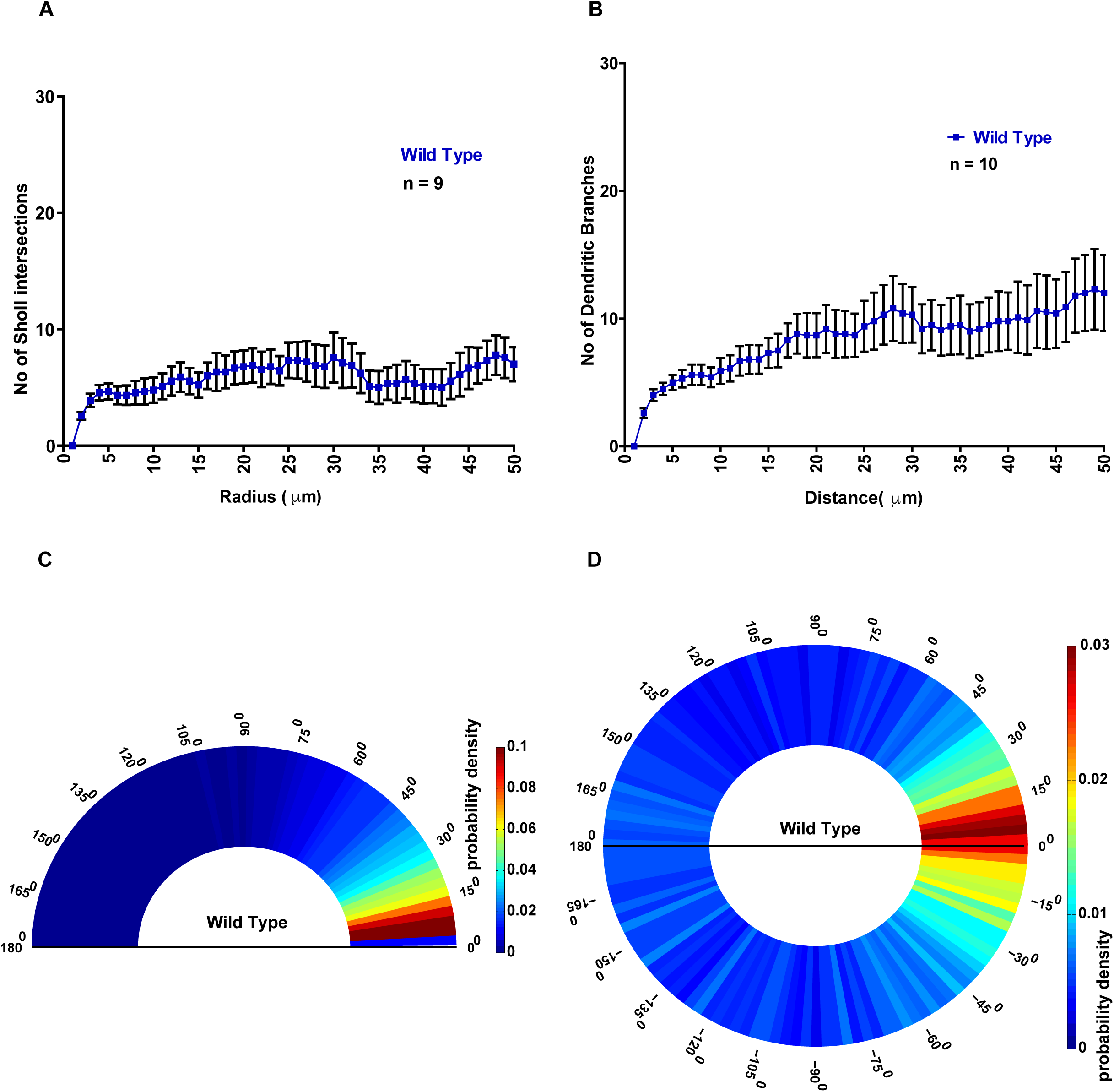
Average number of Sholl intersections, dendritic branches, branching and orientation angle of the apical dendrites in the stratum radiatum of the CA1 region of wild-type hippocampus. (A) The average number of Sholl intersections of pyramidal neurons in the wild type CA1 region of the hippocampus at a distance of 50 μm from the soma. (B) The number of dendritic branches emanating from pyramidal neurons in the CA1 region of the wild type hippocampus. (C) Heat map to analyze the probability distribution of the dendritic branching angle. (D) Heat map to analyze the probability distribution of the orientation angle of CA1 dendrites in wild-type animals. Each wedge or bin represents 2.8 degrees. Error bars represent mean ± SEM.

Basal dendrites of CA1 neurons in the stratum oriens are innervated by recurrent local collaterals and by the CA3 neurons lying closest to CA1 (Ishizuka et al., 1990). The basal dendrites integrate synaptic information very distinctly from the apical dendrites. It has been suggested that the attenuation of EPSPs is steeper in basal dendrites than the apical dendrites when they propagate to the soma (Inoue et al., 2001). The mode of input summation in basal dendrites is location – dependent, in contrast to the spatial amplification of temporally clustered inputs processed by apical or distal dendrites (Nevian et al., 2007). We set forth to understand whether there is a difference in the Sholl intersections and in the orientation angle of basal dendrites compared to that of the apical dendrites of the pyramidal neurons in the CA1 of the hippocampus. Wild type animals exhibited an average Sholl intersection value of 13.53 ± 0.61 (mean ± SEM) for the basal dendrites (Fig. 3A). The branching angle distribution of basal dendrites was similar to that of the apical dendrites (Fig. 3B). Analysis of the relative normalized frequency of basal dendrites in SO (Stratum Oriens) mice showed differential distribution compared to the apical dendrites in the SR (Stratum radiatum). Basal dendrites in the SO showed a strong inclination for dendrite orientation angles ranging from – 180 to -140. They showed a probability distribution of 34.38% for this range of angles. They also exhibited another distribution peak ranging from + 180 to +140 with a normalized relative frequency of 31.2% (Fig.3C). The heat map for the probability distribution of the orientation angle of basal dendrites in the stratum oriens of wild type animals exhibited highest preferred orientation from - 150 to -180 and +150 to +180 degrees with a peak from -160 to -180 and +160 to +180 degrees and a mean probability distribution of 0.008 for the remaining zones (Fig. 3D). Thus high resolution imaging and 3D reconstructions enabled us to discover that the distribution of orientation of the basal dendrites is different from orientation of apical dendrites. We didn’t attempt to extract information on the basal dendrites of pyramidal neurons in *reeler* as some of the apical dendrites are inverted and they are positioned in SO, which renders the analysis difficult to follow. Nevertheless, this approach would be able to distinguish and compare any structural changes of dendrites in other mutants, which are less severe than *reeler* mice to that of the wild type mice.

**Figure 3.**
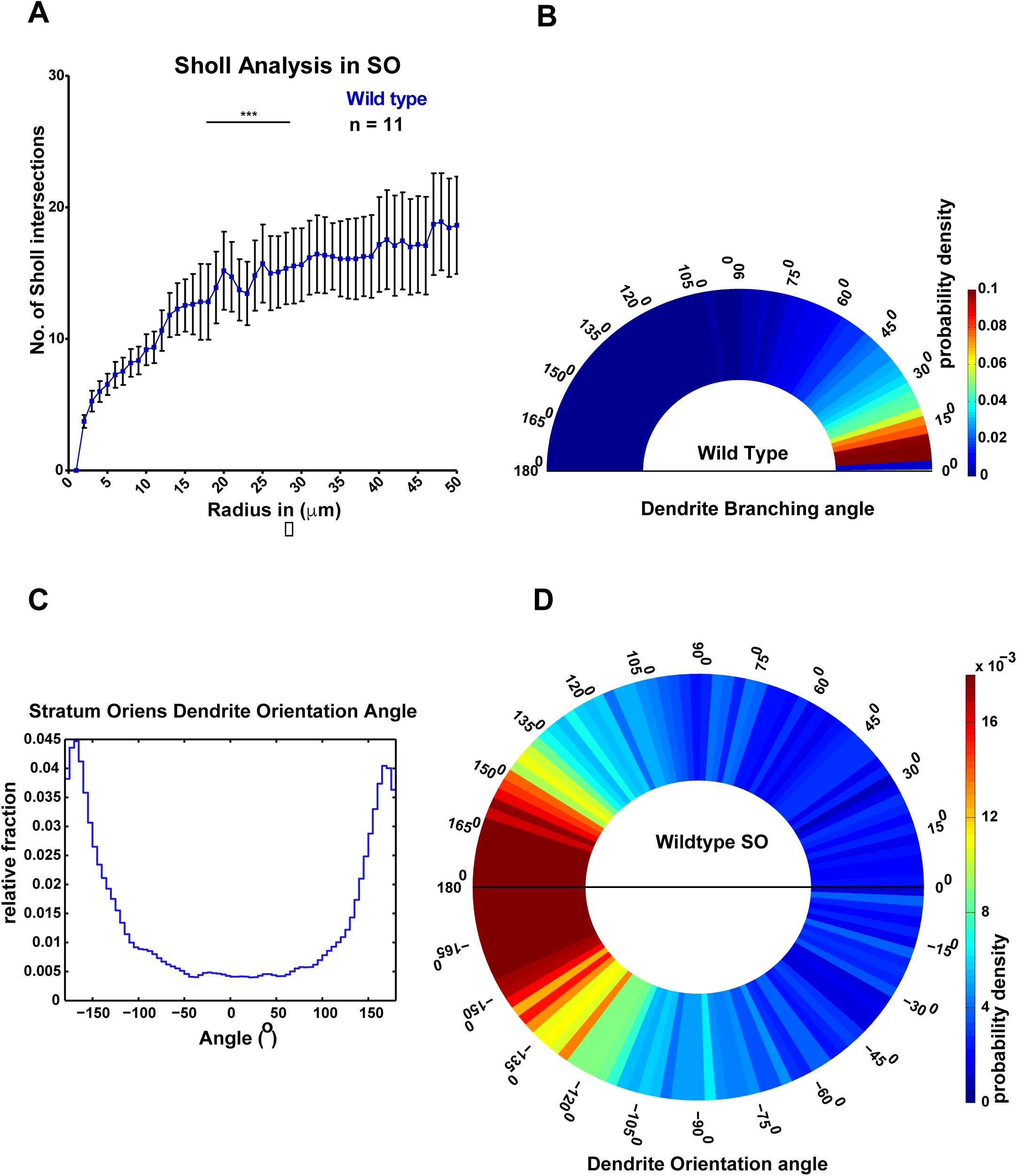
Number of Sholl intersections of the basal dendrites in the stratum oriens of the CA1 region at a distance of 50μm from the soma, dendritic branching angle and dendritic orientation angle in wild type mice. (A) Sholl analysis of the basal dendrites of the CA1 neurons of wild type in the stratum oriens. Error bars represent standard error of the mean (SEM). (B) Heat map denoting the distribution of the branching angle of basal dendrites. (C) Quantitative difference in the distribution of the orientation angle of the basal dendrite in wild type animals. Wild type mice exhibited a preferential inclination of orienting in the angles range from – 180 / +180 to - 140 / + 140 degrees. (D) Heat map depicting the probability distribution of the orientation of basal dendrites in the stratum oriens. Each wedge represents 2.8 degrees.

### 3.2. CA1 apical dendrites of *reeler* are oriented arbitrarily at all angles

Given that the superficial layer is more normal than the deep one of *reeler* CA1, we decided to focus to do the analysis on the superficial pyramidal layer of the *reeler* hippocampus and accordingly defined a ROI (x = 342.67 μm, y = 117.79 μm and z = 40 to 50 μm) that comprised large parts of stratum radiatum in wild type and the apical dendritic region of the superficial pyramidal layer in *reeler*. Apical dendrites in wild-type mice clearly showed preferred orientation angles. In contrast, the *reeler* apical dendrites (apical dendrites of the superficial pyramidal layer) were almost arbitrarily oriented at all angles ranging from - 180 to +180 degrees. In wild-type mice 53.95% of apical dendrites exhibited an orientation preference for the angles ranging from -40 to +40 degrees, whereas the dendrites in *reeler* showed an orientation preference of 31.87% for this range of angles. The remaining apical dendrites, 46% in wild-type mice and 68% in reeler mutants, respectively, were distributed at other angles (Fig. 4A). Furthermore, *reeler* mice showed no inclination towards any preferred range of angles. Instead, dendrites were distributed almost arbitrarily at all angles from -180 to 180 degrees (Fig. 4B). However, the distribution of the branching angles of apical dendrites of both wild-type animals and *reeler* mice showed a similar pattern (Supplementary Fig. 2).

**Figure 4.**
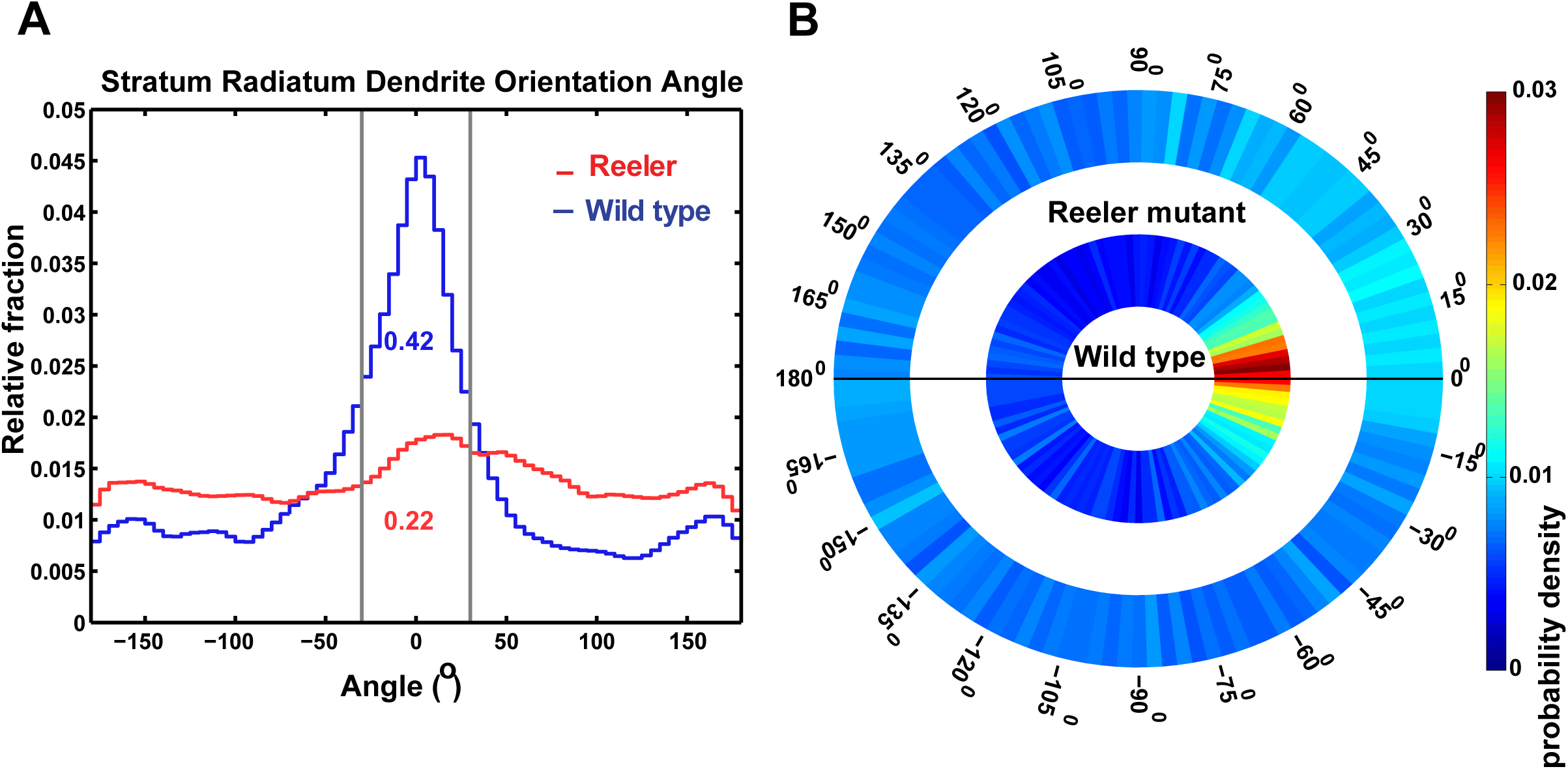
Orientation angle of apical dendrites in the stratum radiatum of *reeler* (apical dendrites arising from the superficial pyramidal layer) and wild-type mice. (A) Quantitative differences in the orientation angle of apical dendrites in *reeler* and wild-type mice. Wild-type apical dendrites exhibited an inclination of preferential orientation from -40 to +40 degrees. In wild-type animals, 42% of the dendrites show angles ranging from -20 to +20 degrees, while in *reeler* mutants 22% of the dendrites expand in this range of angles. (B) Heat map of the probability distribution of apical dendrite orientation in stratum radiatum of wild-type animals and *reeler* mutants (apical dendrites of superficial pyramidal layer).

### 3.3. Quantitative analysis of spine types in SR and SO revealed a significant increase in the number of stubby spines

Dendrite branching pattern, orientation, and development are closely connected with synapse size and maturation (Nimchinsky et al., 2002; Yuste, 2011). Synaptic structural changes such as the size, motility, maturation and potentiation of synapses dictate the plasticity of neurons which is important for learning and memory (Matsuzaki et al., 2004; Sabatini et al., 2001). Spines come in different types such as stubby, thin and mushroom. Reconstruction of the whole CA1 region by stitching imaged tiles enables us to distinguish very clearly between the apical and the basal dendrites emanating from individual pyramidal neurons in wild type mice. Using this approach, one can look into any specific branches of dendrites (apical, distal, and basal) and analyze parameters such as the spine density, types of spines, head and neck volume. In this study, we focused on examining the different types of spines for purposes of methodological validation. To further understand whether there was a difference in the classification of spines in different compartments or regions in CA1, spine classification was performed in the SR and SO region. Morphometric analysis and classification of spines into different groups (Thin, Stubby, and Mushroom) was done in Neuron Studio (http://research.mssm.edu/cnic/tools-ns.html) with reference to the published work (Dumitriu et al., 2011). In the stratum radiatum of CA1 in wild type, pyramidal neurons possessed more stubby spines when compared to mushroom spines. The average number of the relative fraction of stubby spines (normalized) was 0.43 ± 0.021 (mean ± SEM), while that of thin and mushroom spines was 0.29 ± 0.014 and 0.25 ± 0.007 (mean ± SEM) respectively (Figure 5A). In stratum radiatum, the neurons exhibited significantly stubby spines compared to thin and mushroom spines (p < 0.05, one way – ANOVA analysis for multiple comparisons). The basal dendrites in the stratum oriens of wild type mice possess more stubby spines when compared to thin and mushroom spines (p < 0.05, one way – ANOVA analysis for multiple comparisons). The average fraction of stubby spines in the stratum oriens was 0.39 ± 0.014 (mean ± SEM), while that of thin was of 0.32 ± 0.013 (mean ± SEM). The number of thin spines in the basal dendrites of CA1 of the wild type hippocampus was larger than that of thin spines in the apical dendrites (0.29 ± 0.014, Fig 5B). The stubby spines decreased by 4%, while the thin spines increased by 2% in basal dendrites compared to the apical dendrites in the stratum radiatum. The average fraction of mushroom spines in the basal dendrites was of 0.27 ± 0.004 (mean ± SEM) showing an increase of 2% compared to the percentage of mushroom spines in the apical dendrites in the stratum radiatum. This data indicates that in the basal dendrites of CA1, there was an equal increase in the number of thin and mushroom spines compared to the apical dendrites in the stratum radiatum. There was a striking pattern in all the tiles we analyzed, for the majority of the tiles there were more stubby spines compared to other types both in the apical and basal dendrites of the pyramidal neuron in the CA1 region of wild-type, hence the question of oversampling is ruled out (Supplementary Figure 3). These data imply that the CA1 neurons of wild type might be more plastic as there are more stubby and thin spines than mushroom spines (Matsuzaki et al., 2004).

**Figure 5.**
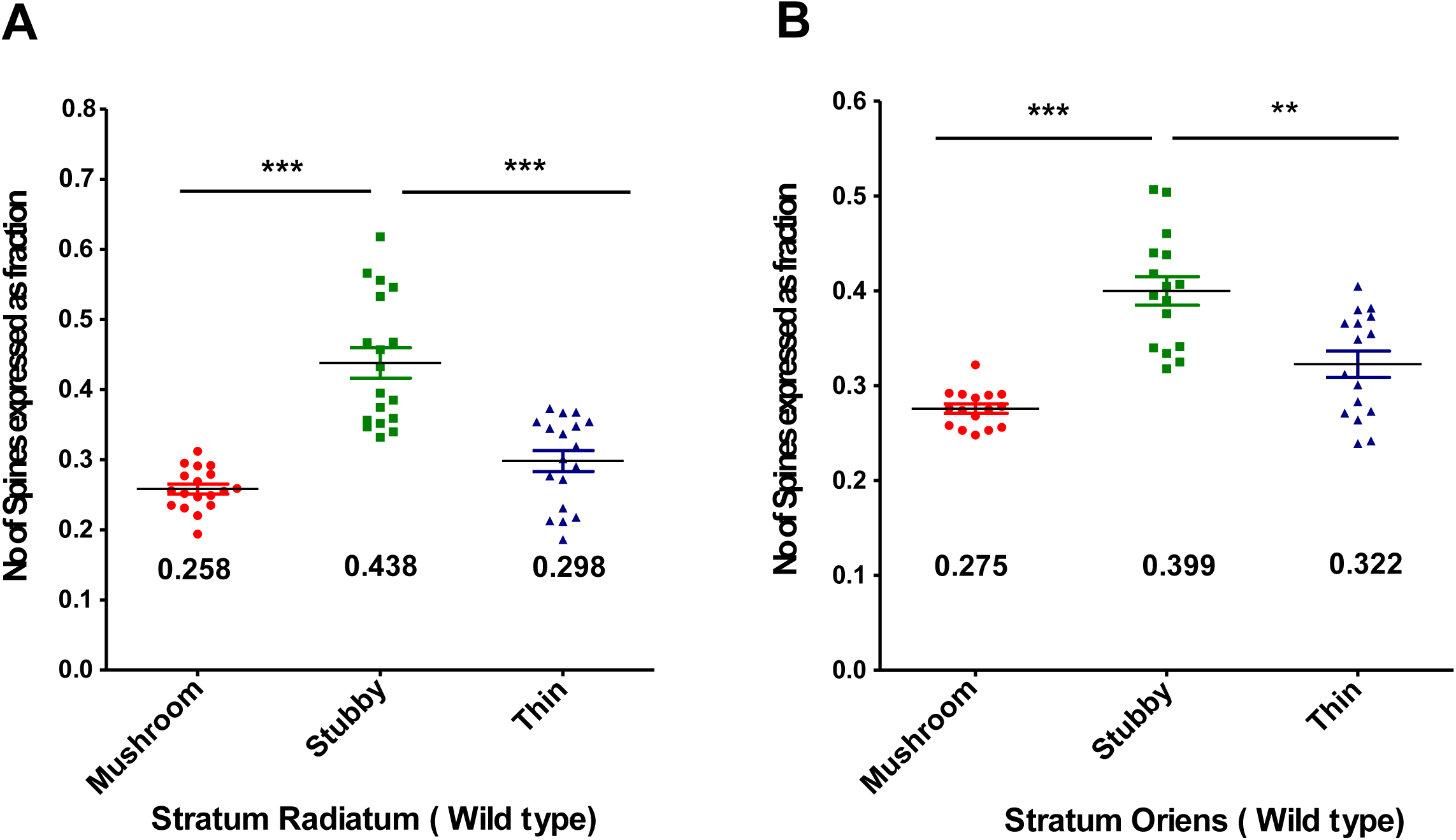
Relative fraction of spine types in the SR, SO in the CA1 region of wild type animals. A) The number of different types of spines in the stratum radiatum of CA1 in wild type. In stratum radiatum, the neurons exhibited significantly stubby spines compared to thin and mushroom (***p < 0.0001, unpaired t-test) B) The distribution of different types of spine in the stratum oriens of the CA1 region of wild type hippocampus. Stubby spines were dominant in the basal dendrites, however, in contrast to the apical dendrites, there were a greater number of thin spines and mushroom spines in the basal dendrites. (***p < 0.0001, **p < 0.001 unpaired t-test).

## 4. Discussion

Using high resolution imaging, we were able to systematically characterize structural properties of the pyramidal neurons in the CA1 region of hippocampus. We quantitatively measured the number of Sholl intersections, dendritic branches, the branching and orientation of dendrites as well as the types of spines in the CA1 region of wild type hippocampus. Using tile based imaging of the whole region of the CA1 aided by a custom made macro enables us not only to reconstruct the whole CA1 region, but also provides a unique opportunity to extract key anatomical features which might be useful to understand the network properties of the circuits these neurons are a part of. One of the potential uses of this method is to compare the structural properties of the wild type neurons to that of the neurons which are defective for a protein of interest. To substantiate this claim, we compared the apical dendrites emanating from CA1 neurons residing in the superficial layer of the reeler mutant hippocampus to that of the apical dendrites from the pyramidal neurons in the CA1 region of wild type hippocampus.

The geometry of the dendritic arborizations plays an important role in the synaptic integration of inputs. This will have a profound effect on the excitability of dendrites, their integrative abilities such as the association, co-operation and integration of different synaptic inputs as well as on the generation of dendritic spikes, which enable long-term potentiation in activated synapses. Using high resolution imaging, we could quantify the number of Sholl intersections, the branching and orientation of the apical and the basal dendrites and the different types of spines present on them. Given that the apical dendrites integrate information differently from the basal dendrites, quantifying the structural parameters such as the number of intersections and branches will give insights on the complexity of the network the neurons are part of. Utilizing this approach, one can also image the axon arborization to accurately measure the connectivity matrix of the pyramidal neurons. Surprisingly we found that the orientations of apical and basal dendrites are different. 53.95% of apical dendrites exhibited an orientation preference for the angles ranging from - 40 to +40 degrees, the remaining apical dendrites, (46%) were distributed at the remaining angles while the basal dendrites showed a strong inclination of orienting their dendrites at the angles ranging from – 180 to -140 with a probability distribution of 34.38% for this range of angles. The basal dendrites also showed another distribution peak ranging from + 180 to +140 with a normalized relative frequency of 31.2%. The functional significance of this preferential distribution of dendrites is unknown, however mapping the axon arborization data to determine the connectivity matrix might yield a functional implication. We also showed that both the apical and basal dendrites possess more stubby spines than mushroom ones. The functional significance of the increase in the number of stubby spine is unknown.

### 4.1. The connectivity matrix is altered in the *reeler* hippocampus

We imaged the apical dendrites of the superficial layer of CA1 in reeler hippocampus to unravel the anatomical changes exhibited by this mutant. Morphological analysis of the apical dendrites in the CA1 neurons in wild-type and *reeler* mice showed striking differences in the geometry of the dendritic branching pattern and this might change the connectivity matrix. Pyramidal neurons in *reeler* mice show an orientation of apical dendrites at all angles ranging from -180 to +180 degrees in a rather arbitrary way in contrast to the preferential orientation of apical dendrites in wild-type animals. This in turn might affect the connectivity matrix of the *reeler* hippocampus. The dendritic orientation of *reeler* mice might facilitate randomly formed synaptic connections and multiple hits within the same dendritic segment. The findings of the present study suggest there might be a perturbed connectivity matrix of CA1 pyramidal neurons in *reele*r mice. The dendrites in *reeler* are more branched but shorter. This could alter their synaptic connectivity with entorhinal fibers which (also in *reeler* mutants) terminate on distal dendritic segments (Zhao et al., 2003). Because there are an increased number of short branches in the proximal region of the dendritic tree in stratum radiatum, the number of proximal (commissural/associational) synapses is likely to be higher in *reeler*. Taken together, this would amount to a sparser and weakly connected network in *reeler* mice. The major implication of this result is that *reeler* mice might have a perturbed connectivity matrix and hence different network properties. Knowing how many active inputs impinge on these morphologically altered dendrites and spines is a very important issue to be addressed. What are the functional properties of the synapses in this matrix? Besides that, it is very important to know the axon arborization to determine accurately the connectivity matrix in *reeler* mice to understand its functional implications. Given that not all geometrically apposed synapses are functional, further work needs to be done to understand how the perturbed orientation of dendrites and spines in *reeler* affects the functional properties of the neurons, and further of the network.

### 4.2. The distribution of different spine types in the CA1 of wild type hippocampus

We utilized an imaging approach to unravel whether there is a difference in the distribution of different types of spines on the apical and basal dendrites of the CA1 region. In this study, we restricted our analysis only classifying different types of spines; however, one can also measure the spine density, spine head volume and spine neck length. This study demonstrates that wild type mice (3 months of age) have more stubby spines compared to thin and mushroom. Mushroom spines are considered to be the spines that store traces of memories, while the small spines such as the stubby are the “learning spines” undergoing synaptic plasticity (Matsuzaki et al., 2004). It has been speculated that enhanced synaptic transmission can convert thin spines and stubby spines (learning spines) into mushroom spines (memory spines, Matsuzaki et al, 2004). As there is no direct physiological correlation for an increase in particular types of spines, which can be translated into the behavioral ability of performing a task in animals, the increase in stubby spines does not necessarily imply an increase in cognitive abilities. Generalizing from such facts needs to be done very cautiously as spines is not static and can change form. Even the thin protrusions make synapses, and there is no absolute correlation between the stability of the spines and their size. Moreover, LTP induction protocols and electrophysiological recordings in slices have shown that LTP was induced in the slices; however, the spines which were potentiated could not be identified (Lang et al., 2004), indicating that changes in the morphology of spines do not necessarily indicate the induction or the expression of synaptic plasticity. The present study demonstrating morphological changes in dendrite organization in association with changes in spine size thus clearly points to functional implications. In essence, this type of large scale anatomical study can be done on any animals harboring a particular mutation and even subtle changes in the dendritic architecture can be dissected by high resolution imaging approaches. The anatomical details gleaned from this study might help in asking specific physiological questions to understand functional network.

## Figure legends

**Supplementary Figure 1:**
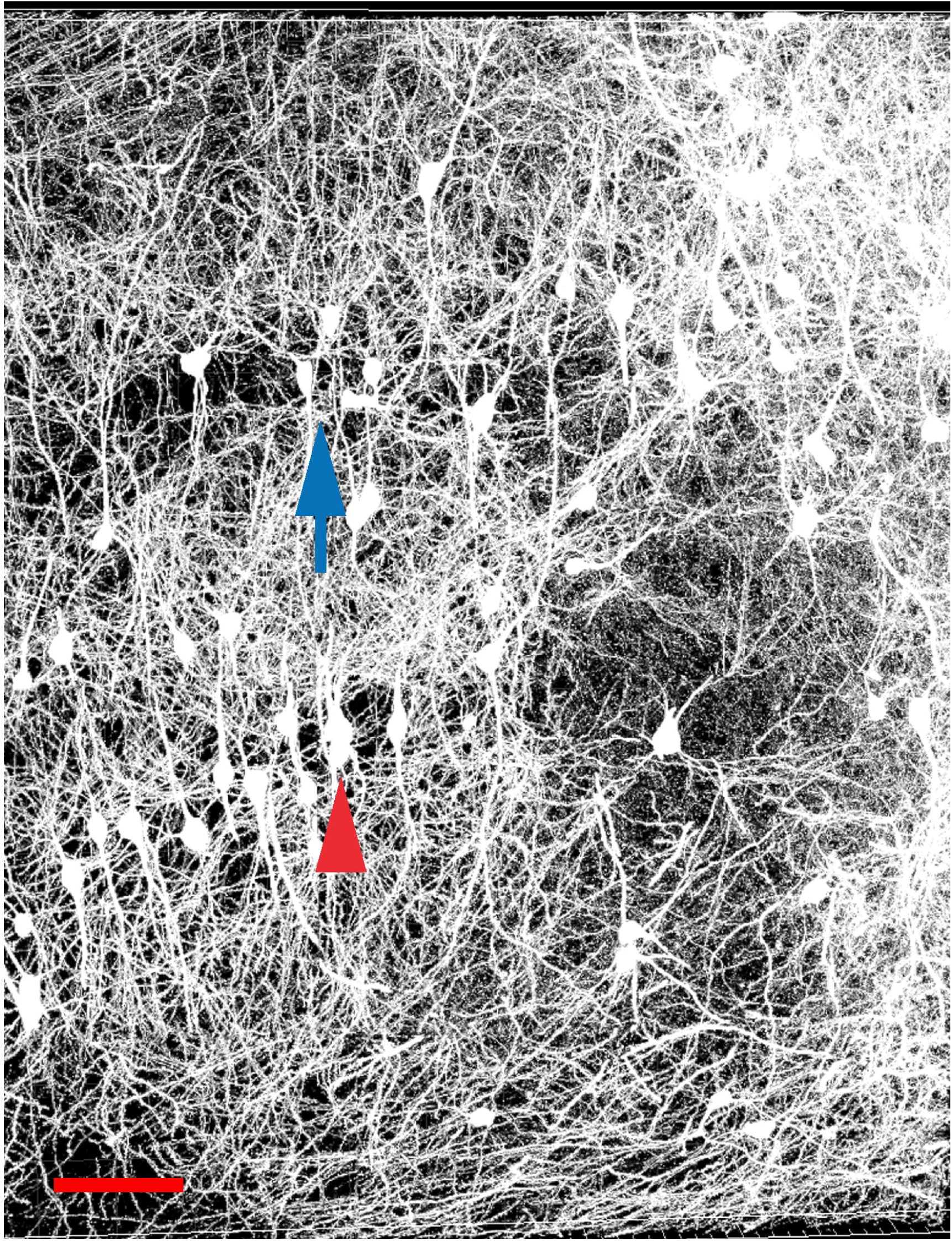
Reconstruction of the whole CA1 region of *reeler* hippocampus. The CA1 region of *reeler* mouse hippocampus. The pyramidal layer is split into two: a deep pyramidal layer (Blue arrow) and a superficial pyramidal layer (Red arrow). The dendrites display inverted orientation and are misplaced. All dendrites in the CA1 region are reconstructed using Filament tracer plug – in of Imaris bitplane. Scale bar: 70μm.

**Supplementary Figure 2:**
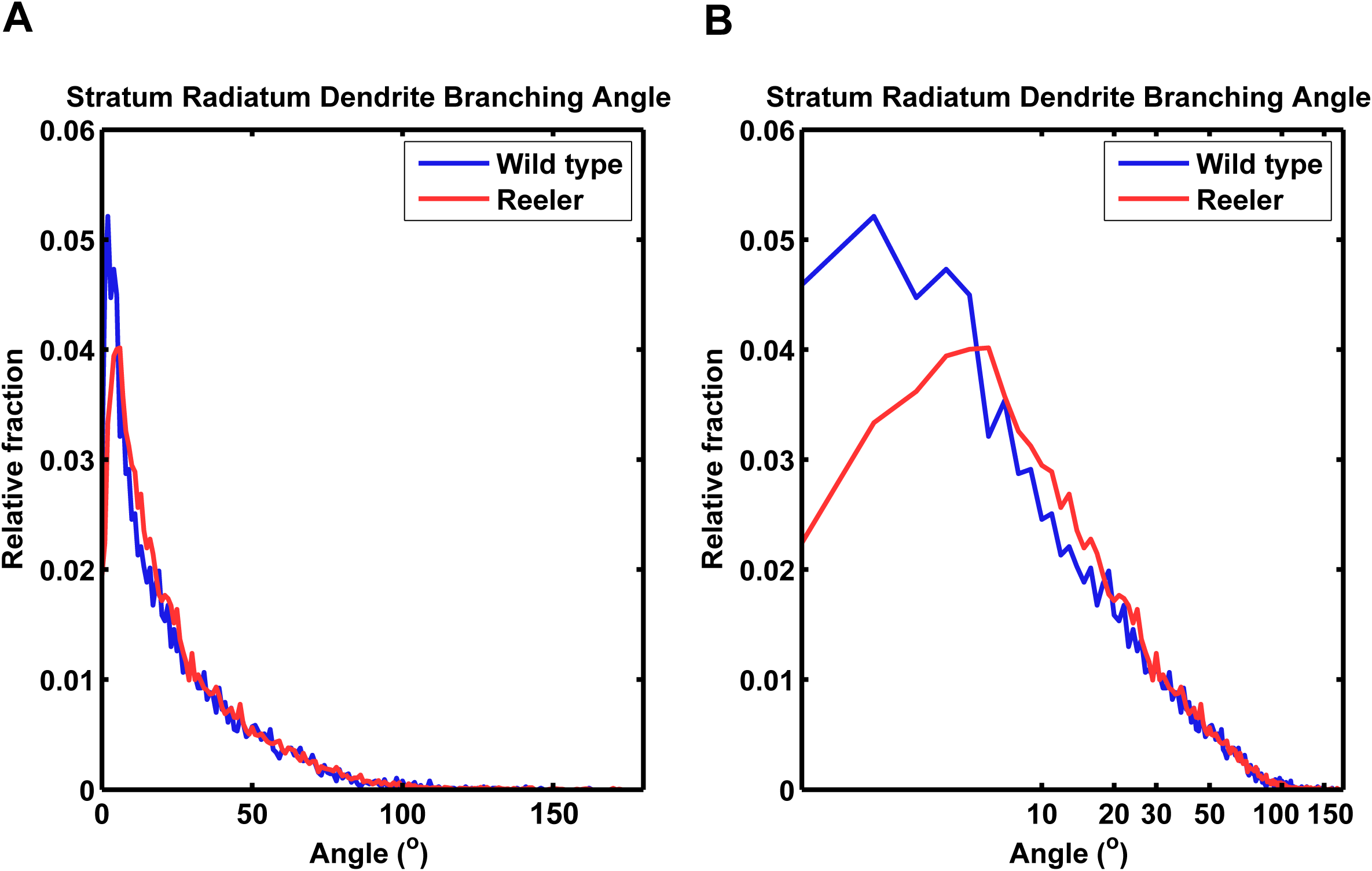
Quantitative analysis revealed no significant difference in the dendritic branching angle of apical dendrites in *reeler* and wild type. Relative fraction of the frequency distribution of the dendrite branching angle of the CA1 neurons of stratum radiatum in wild type and *reeler* mice. No statistical significance was observed between *reeler* and wild type data. To better highlight the loss of any significant difference in the branching angle, the relative fraction of the branching angle was plotted on a log scale. A) In Left panel, branching angles were presented in a natural scale. B) In Right panel, branching angles were presented in logarithmic scale.

**Supplementary Figure 3.**
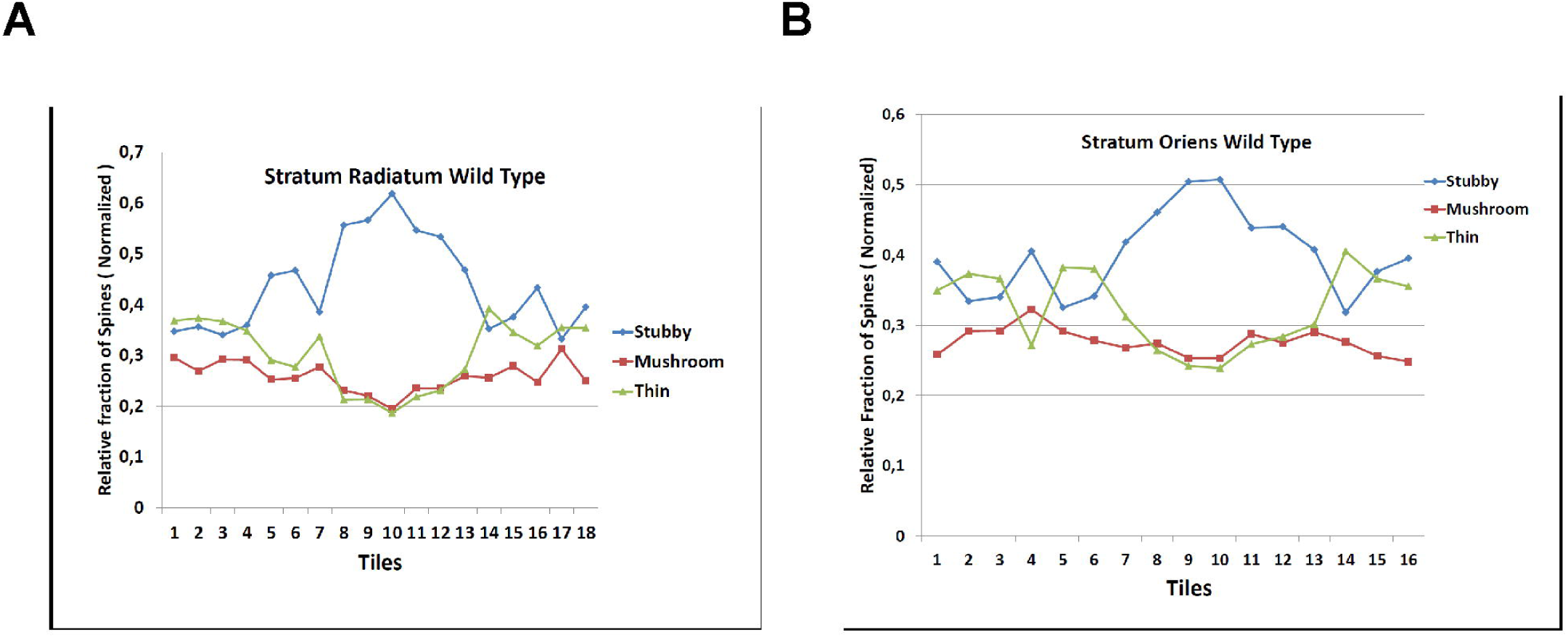
Analysis of the difference in the number of different types of spines in individual tiles in the CA1 region of hippocampus of wild type mice. A) The relative fraction of spines in each tile in the stratum radiatum of the CA1 region of the hippocampus. B) The relative fraction of spines in the stratum oriens of wild type mice in each tile. Note that in most of the tiles, stubby spines are predominated.

## Acknowledgements

We acknowledge Dr. Shaojun Jin for providing the macro to image the hippocampus and Dr. Roland Nitschke for his assistance in imaging in the Life Imaging Center of the Center for Biosystem Analysis at the University of Freiburg, Germany. We are grateful to Dr. Arvind Kumar (Bernstein Center for Computational Neuroscience, Freiburg) for his critical comments on this work and for many valuable discussions. The work presented in this study was performed in partial fulfillment of the requirements for a PhD at the Faculty of Biology, University of Freiburg, Germany (M.Y.) and was supported by the Deutsche Forschungsgemeinschaft (FR 620/12-1). MF was a Senior Research Professor supported by the Hertie Foundation. <u>MY dedicates this work to his PhD mentor Late Dr. Michael Frotscher (MF).</u>

## Abbreviations

EPSPs: Excitatory post-synaptic potentials
APs: Action potentials
CA1: *Cornu Ammonis*

